# Method for *in vivo* assessment of cancer tissue inhomogeneity and accurate histology-like morphological segmentation based on Optical Coherence Elastography

**DOI:** 10.1101/2020.02.06.937417

**Authors:** Anton A. Plekhanov, Marina A. Sirotkina, Alexander A. Sovetsky, Ekaterina V. Gubarkova, Sergey S. Kuznetsov, Alexander L. Matveyev, Lev A. Matveev, Elena V. Zagaynova, Natalia D. Gladkova, Vladimir Y. Zaitsev

## Abstract

We present a non-invasive method based on Optical Coherence Elastography (OCE) enabling the *in vivo* segmentation of morphological tissue constituents, in particular, monitoring of morphological alterations during both tumor development and its response to therapies. The method uses compressional OCE to reconstruct tissue stiffness map as the first step. Then the OCE-image is divided into regions, for which the Young’s modulus (stiffness) falls in specific ranges corresponding to the morphological constituents to be discriminated. These stiffness ranges (characteristic “stiffness spectra”) are initially determined by careful comparison of the “gold-standard” histological data and the OCE-based stiffness map for the corresponding tissue regions. After such precalibration, the results of morphological segmentation of OCE-images demonstrate a striking correlation with the histological results in terms of percentage of the segmented zones. To demonstrate high sensitivity of the OCE-method and its excellent correlation with conventional histological segmentation we present results obtained *in vivo* on a murine model of breast cancer in comparative experimental study of the efficacy of two anti-tumor chemotherapeutic drugs with different mechanisms of action. The new technique allowed *in vivo* monitoring and quantitative segmentation of (i) viable, (ii) dystrophic, (iii) necrotic tumor cells and (iv) edema zones very similar to morphological segmentation of histological images. Numerous applications in other experimental/clinical areas requiring rapid, nearly real-time, quantitative assessment of tissue structure can be foreseen.

## Introduction

Histopathology still remains the most reliable “gold standard” method for assessing the histological changes. However, this method is invasive, rather time-consuming and labor-intensive, requiring both the preparation of histological sections (including staining) and their interpretation/assessment. The histological image assessment and segmentation is performed by a histopathologist, which makes the results operator-dependent. Furthermore, the invasive/destructive nature of the histological examinations, in principle, makes it impossible to perform *in vivo* non-invasive monitoring of the tissue changes. In addition, taking a biopsy may not fully reflect the overall picture of the tumor, due to its heterogeneity.

In view of this, significant attention has been paid to the development of alternative fast and non-invasive ways of obtaining information about biological tissues similar to the results of histological image interpretation. In this context, various optical methods have been proposed, in particular the use of variations in auto-fluorescence properties to detect tumor cells**^1,2^** or the generation of higher harmonics, which can be used to detect the presence of collagen and elastin**^3^**. The use of Optical Coherence Tomography (OCT) for such purposes is also attracting much attention, because the characteristic scales of OCT scans occupy an intermediate place between macroscopic imaging (like medical ultrasound) and cell-resolution microscopy. OCT scans typically provide several millimeters lateral field of view to a depth of ~ 1-2 mm, and these are comparable with the ranges of histological images. While the intrinsic resolutions of OCT (typically 5-10 μm) and histology (typically cell-level) may differ depending on the specific implementation of the former, the spatial scales at which tissue structures and types are delineated and segmented (e.g., tumor vs normal, viable vs necrotic regions, etc.) are similar for the two techniques. Since different tissue components have different optical properties, the manifestations of such differences in OCT scans can be used to perform such segmentation/delineation. For example, differences in birefringence can be used for the detection of collagenous components**^4^** or for distinguishing tumor/non-tumor regions during neurosurgery**^5^** by analyzing polarization-sensitive OCT images. For similar purposes, multiparameter analysis of the speckle-pattern features in OCT scans (for example, the statistical properties of speckles, polarization features and signal decay, etc.) can be used (see, e.g.**^6^**). Multimodal OCT has also proven to be very useful for *in vivo* evaluation of fairly rapid and pronounced post-therapeutic changes in tumors. For example, one can mention the OCT-based angiographic observation of blood-circulation blockages in tumors and in peri-tumorous regions for accurate prediction of the outcome of vasculature-targeted photodynamic therapy (PDT) during the first 24 hours post-PDT^6–8^. However, unlike the fairly easily observed perturbation in the microcirculation of blood, the assessment of the histological tissue structure lacks precise, non-invasive methods. In this context, the proposed method of tissue assessment/segmentation using OCE opens unprecedented prospects for such finer quantitative assessment of quite minor changes in the tissue.

In a broad sense, the OCE-approach discussed here is also based on OCT-scan analysis, but in an essentially different way. Namely, the OCT imaging is used merely as an auxiliary step to obtain elastographic maps (i.e., maps of stiffness - the Young’s modulus of the tissue) by applying quantitative phase-sensitive quasistatic compressional Optical Coherence Elastography (OCE)**^9–11^**, and using its realization as described in**^12–15^**. Another key feature of the described method is the application of procedures that can be called “elasto-spectroscopy”**^16^**, the term having been introduced by analogy with “mass-spectroscopy”**^17^** and being based on assessments of certain specific ranges of stiffness within the reconstructed OCE images. It is known that the Young’s modulus of biological tissues may vary by several orders of magnitude. In particular, as various types of tumor are typically stiffer than the surrounding tissues, palpation has been used for centuries to detect tumors as stiffer inclusions embedded in the tissues. With the appearance of quantitative elastography in the 1990s (including such methods as ultrasound- and MRI-based elastography that have become routinely used in clinical situations**^18^**) the possibilities for even more detailed assessment of cancer types have been studied, based on tumor stiffness as estimated with averaging over volumes at ~mm scales or larger, typical of the MRI and US methods**^19^**. The appearance in recent years of new elastographic methods with even higher resolution (first of all, OCE) has demonstrated that tumors, themselves, are rather heterogeneous and multi-component in terms of their elasticity**^16,20–22^**. This OCE-based observation agrees well with histological conclusions that tumor structure comprises various histological components for which different elastic properties can be expected. The proportions of those components strongly depend on the molecular-biological type of the tumor, the stage of tumor development, the applied therapies, etc.

The resolution of OCT-based elastographic methods allows one to resolve stiffness heterogeneities on essentially sub-millimeter spatial scales for both naturally occurring human tumors (e.g., breast cancer)**^16,20^** and model tumors studied in animal experiments**^22^**. As shown in**^16^** for patients’ breast-cancer samples, and will be even more clearly demonstrated here using model tumors with simpler histology, careful comparison between OCE images and histological images of the same tumor zones reveals striking correlations between tissue components and their stiffness. Therefore, quantitative OCE mapping of tumor stiffness with a resolution of several tens of microns opens possibilities to resolve quite fine tissue heterogeneities in OCE images and perform morphological segmentation non-invasively. In the following discussion, it is shown that both methods provide very similar accuracy and resolution of the resultant morphological maps, demonstrating striking correlation between the areas segmented by OCE and those deduced from conventional assessment of the corresponding histology. However, in contrast to time-consuming conventional histology, the proposed OCE-based automated morphological segmentation is non-invasive, applicable *in vivo,* can be performed at several-minute intervals and is much less labor-intensive in comparison with the conventional procedures of fixation, cutting and staining of histological sections and their interpretation by histopathologist.

Such OCE-based morphological segmentation can potentially be used in a wide range of biomedical applications, in which the variations in the state of the tissue are conventionally assessed using examination of histological sections. These applications comprise (but are not limited to) evaluation of the results of photodynamic, radiation or other anti-tumor therapies and monitoring of the natural development of tumor and non-tumor tissues. The richness of information provided by the OCE-based morphological segmentation about the tissue state/structure is comparable with the informative value of conventional histological segmentation. For instructive illustration of the method’s capabilities, we present the results of *in vivo* comparative experimental studies of tumor response to anti-tumor chemotherapy. The method demonstrates high sensitivity to histological structure with the ability to perform previously unavailable rapid, *in vivo* quantitative assessment/segmentation of morphological alterations in tumors in the treated and control groups. Detailed comparison between the proposed OCE-based “elasticity-spectrum” approach and conventional histological examination was performed and demonstrated excellent consistency.

### Justification of the choice of biomedical demonstration

The potential and efficiency of the new diagnostic method can be convincingly demonstrated using an animal tumor model, which is (a) accessible for *in vivo* monitoring by OCE, (b) is readily controllable and (c) is characterized by at least several co-existing evolving tissue constituents, variations in which can be produced by the test procedures (in our case under the action of two chemotherapy drugs).

In this context, the choice of chemotherapy is based on the fact that this type of anti-tumor therapy remains one of the most widely used methods of cancer treatment**^23^**. The efficacy of conventional chemotherapy is far from satisfactory, this being related to the broad variability of properties of tumors and, therefore, their multiple drug-resistance**^24^**. Thus, control of the effectiveness of chemotherapy is important for the disease prognosis, the choice of the treatment tactics (choice of drugs) and the possible recommendation of surgery**^25,26^**. Assessment of chemotherapy efficiency is usually based on evaluation of tumor dimensions (RECIST - response evaluation criteria in solid tumors) measured by MRI or computed tomography. However, these criteria do not reflect other histologic/functional changes that may occur during the tumor evolution.

The development of tumors may be often accompanied by inflammation causing edema. Highly-aggressive tumors with very fast growth often have regions of spontaneous necrosis**^27^**. The destruction of tumor cells by chemotherapy also mainly has the form of necrosis**^28^**. Tumors may also have clusters of dystrophic cells. Therefore, assessment of the above four tumor constituents, namely, (i) clusters of viable tumor cells, (ii) dystrophic tumor cells, (iii) edema and (iv) necrotic tumor cells are of key importance from the viewpoint of evaluation of the state of a tumor and its reaction to the applied chemotherapy. There is currently no other means available than histological examination for the assessment of such morphological alterations in the tissue.

In this study, the ability of OCE to visualize and quantify these morphological alterations in tumors was studied for two anti-tumor chemotherapeutic agents with essentially different mechanisms of anti-tumor action - anti-angiogenic Bevacizumab**^29^**, and Cisplatin, with a direct cytotoxic action**^30^**. Thus, it is important to compare the destructive action of these two principally different drugs by evaluating the corresponding morphological alterations in the treated tumors.

For this purpose the proposed OCE-based approach based on stiffness-spectrum assessment is applied**^16^**. The high efficiently of OCE for the *in vivo* quantitative assessment of differences in the histological reaction of the tumor to chemotherapy by the two drugs is demonstrated.

## Results

### Conventional histological assessment of tumor response to chemotherapy

A macroscopic evaluation of the state of the tumors showed that Cisplatin and Bevacizumab have moderate antitumor activity against the mouse 4T1 breast cancer tumor. The Tumor Growth Inhibition (TGI, see **Eq. (2)** in Material and Methods) coefficient was ~63% for Bevacizumab-treated tumors and about 75% in the Cisplatin-treated group (see **Fig. 1b**). However, the tumors of these groups did not significantly differ, statistically, in volume. Thus, the standard macroscopic evaluation of the antitumor efficacy of chemotherapy drugs in the inhibition of tumor growth did not allow us to identify appreciable differences between the two agents with different mechanisms of action.

**Fig. 1.**
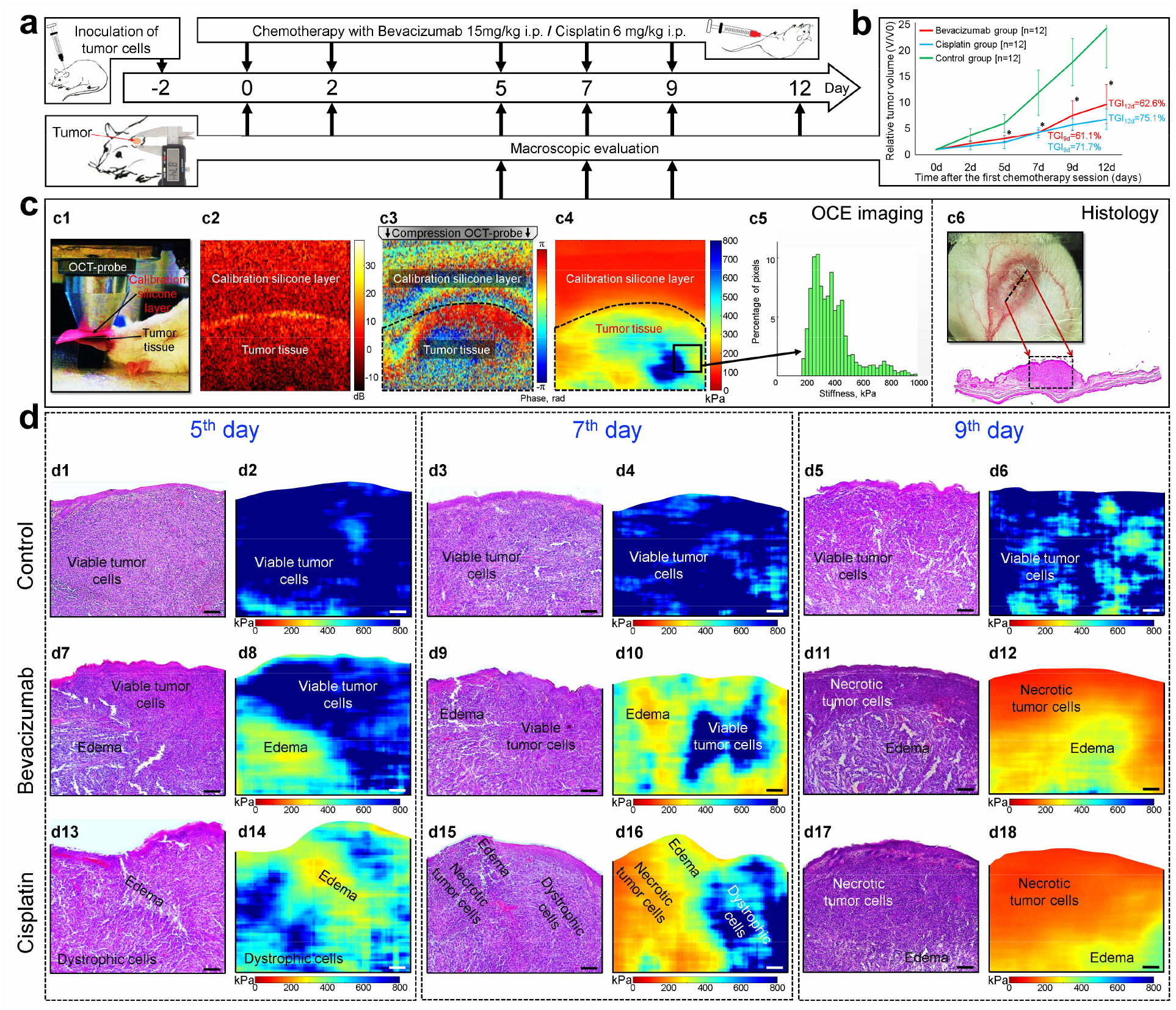
Experimental design, schematically shown OCE-procedures and preliminary results of OCE/histology comparison. **a**, Experimental time course. Two days before treatment started (day −2) tumor cells were inoculated into the ear skin. Chemotherapy was started on day 0 and continued on days 2, 5, 7 and 9. For days 0 to 12 tumor volume changes were visually monitored macroscopically using repeated caliper measurements. OCE imaging was carried out on days 5, 7 and 9. Tumor resection for histological assessment was performed at the OCE time points in three animals to validate the *in vivo* observations. **b**, Kinetics of tumor growth under chemotherapy with Bevacizumab and Cisplatin; the arrows indicate the time point of drug administration; * statistically significant differences from the control group (p<0.05) noted in the Bevacizumab and Cisplatin groups. **c**, Elucidation of experimental OCE procedures for imaging of tumor. OCT probe pressing onto the studied sample, structural OCT B-scan of cancerous tissue under reference silicone, inter-frame phase variation, OCE image and histogram showing the stiffness spectrum (histogram showing percentage of pixels with a particular stiffness value) over a selected region of the OCE image. **d**, Representative examples of parallel monitoring of tumor evolution during chemotherapy by conventional histological examination (left columns) and the stiffness maps obtained by OCE method (right columns) that will be segmented as explained below. Scale bar for all images = 100 μm.

By contrast, histological analysis of tumor sections stained by hematoxylin and eosin revealed distinctive features of the action of the two chemotherapeutic agents, corresponding to specific morphological alterations in the treated tumors. On day 5 (see experimental design **Fig. 1a**), tumors of the control group consist of closely spaced, viable tumor cells that occupied over 90% of the histological image area (see **Fig. 1 d1**). In Bevacizumab-treated tumors, in addition to viable tumor cells, extensive areas of pronounced edema were revealed on day 5, occupying up to 47% (see **Fig. 1 d7**). For Cisplatin-treated tumors, along with viable tumor cells, areas of dystrophic cells (irreversibly damaged) and edema, respectively occupying 45% and 20%, were found (see **Fig. 1 d13**).

On day 7 the appearance of regions of dystrophic tumor cells (13%) and small zones of weak edema (3%) were observed in the control group (**Fig. 1 d3**). In Bevacizumab-treated tumors, the zones of pronounced edema occupied over 63% (**Fig. 1 d9**). In Cisplatin-treated tumors in addition to zones of dystrophic tumor cells, necrotic tumor cells zones (27%) were appeared (**Fig. 1 d15**).

Finally, on day 9 the necrotic tumor cells zone was significantly larger in Cisplatin vs Bevacizumab group and amounted to 69% and 20%, respectively (see **Fig. 1 d11, d17**).

Thus, the histology for all three groups consistently indicates the presence of the following main morphological alterations that arise during chemotherapy: (1) necrotic tumor cells, which occurs in both treated groups, but mainly in the Cisplatin-treated group, due to the specific action of this drug; (2) dystrophic tumor cells were found in all the groups, since this state precedes tumor cell necrosis; (3) edema, which was detected in all groups as a result of tumor development, however, it was the most pronounced in the Bevacizumab-treated group, being caused by the specific (targeted) damage to the vessels walls. The initial constituent (4), viable tumor cells, was also found in all three studied groups, however, it dominated only in the control group (compare the histological images for the three groups in **Fig. 1d**).

### Obtaining elastographic stiffness maps and their targeted comparison with histological images

The main physical principles of the realization of compressional OCE were described in**^11–13,31^** and are summarized in the OCE-section of Materials and Methods. Schematically the essence of OCE-examinations is illustrated in **Fig. 1c**. These panels show that the key step in OCE is the quantitative mapping of mechanical strains in the examined tissue subjected to mild (of the order of a few percent) mechanical compression by the OCT-probe (shown in **Fig. 1c1**) through a translucent layer of silicone with a pre-calibrated Young’s modulus (see structural OCT image in **Fig. 1c2**). The axial gradients of interframe phase variations (shown in **Fig. 1c3**) can be recalculated into local strains over the OCT frame and then comparison with strains in the reference layer makes it possible to convert the relative strain distribution into a quantified map of the Young’s modulus (stiffness) of the tissue as shown in **Fig. 1c4**. Of key importance is that a single stiffness map as that shown in **Fig. 1c4** is actually generated from several tens of raw OCT images obtained during the tissue compression, such that every vertical column in the resulting stiffness map corresponds to the *same* pre-chosen pressure exerted by the silicone layer on the tissue. The point is that when acquiring individual OCT images, the stress applied by the silicone to the tissue is usually essentially inhomogeneous over the frame because the tissue boundary is normally curved and the tissue structure is mechanically very heterogeneous. This geometrical/structural inhomogeneity results in significantly different loading of the tissue, which may strongly (e.g., by a factor of several times) affect the ‘measured’ tissue stiffness because of the pronounced nonlinearity of the ‘stress-strain’ dependences for real tissues**^15,16,32^**, meaning that it makes no sense to interpret the tissue stiffness without specifying (or standardizing) the loading conditions. Thus, the performed pressure standardization in the synthesized OCE scan (see OCE-section of Materials and Methods) ensures meaningful interpretation of the tissue stiffness over the entire OCE scan, independently of the tissue boundary shape and internal heterogeneities.

Pair-wise examination of the resulting stiffness maps with the corresponding histological sections similar to those shown in **Fig. 1d** clearly demonstrate that the morphologically differing tumor zones revealed in the histological images do correlate very well with the stiffness variations.

Thus, on day 5 of observation the control tumors represented on the OCE images were characterized mainly by maximal stiffness values (see **Fig. 1 d2**) that correspond to densely packed viable tumor cells. On days 7 and 9, the appearance of softer areas on the OCE images was observed, and these softer ‘patches’ probably correspond to zones of dystrophic tumor cells and zones of weak edema and occupy 10-20% of the OCE image (see **Fig. 1 d4, d6**).

By contrast, in the therapeutic groups the stiffness variations are much more pronounced. For Bevacizumab-treated tumors, by day 5 of observation, the appearance of larger areas with reduced stiffness values (see **Fig. 1 d8**) is already characteristic, these corresponding to edema on the histological images. On day 7 of observation, areas with reduced stiffness values (greenyellow color) prevail over areas with high stiffness values (dark-blue color) on the OCE images (**Fig. 1 d10**) and these topographically coincide with zones of pronounced edema in the histological images. The blue-color regions with high stiffness values that persist in **Fig. 1 d10** coincide with the zones of viable tumor cells in the histological images **Fig. 1 d9**. On day 9, there was a further decrease in stiffness, so that the most stiff blue-color areas disappeared, whereas an area with even stronger-reduced stiffness values near the tumor surface appeared (see the orange-color regions in **Fig. 1 d12**) that topographically coincided with necrotic tumor cells zone on the histological images.

For Cisplatin-treated tumors, by day 5, it was also observed the appearance of large areas with somewhat reduced stiffness value (green-yellow color in **Fig. 1 d14**), corresponding to zones of dystrophic tumor cells and edema in the histological images. On day 7 there were noted: (i) regions with reduced stiffness values (light-blue and greenish colors) consistent with zones of dystrophic tumor cells on histological images; (ii) areas with even lower stiffness values (yellow-green color in the OCE-map) consistent with zones of edema in the histological image; and (iii) areas with very low stiffness values (orange-color regions), topographically consistent with necrotic tumor cells zone (see **Fig. 1 d16**). OCE images obtained on day 9 were characterized by a predominance of orange-color areas of low stiffness values over yellow-green areas of moderately reduced stiffness values (see **Fig. 1 d18**). These areas in the stiffness maps were topographically consistent with zones of necrotic tumor cells and edema on the histological images.

Thus, comparison between the OCE and the histological images clearly showed that the gradually evolving heterogeneities in the stiffness distribution are closely related to the morphological alterations seen in the histological sections. In what follows we show that deeper comparative analysis of the histological and OCE images makes it possible to use the stiffness maps for quantitative characterization of the morphological constituents of the tumor and detection of the boundaries between the corresponding zones, such that elastographic images can be automatically converted into morphologically segmented images similar to histological images manually segmented by a histopathologist.

### Determining the specific stiffness ranges of the tumor constituents and OCE-image segmentation

The next step in the study was the quantitative processing of the OCE images, from which the characteristic stiffness ranges corresponding to the zones of the main morphological constituents of the tumor could be determined.

We compared the areas of the selected tumor zones corresponding to the main four morphological constituents (**Fig. 2a**) on histological images and the OCE images (**Fig. 2b,c**) previously obtained *in vivo* at the same locations. For illustration, in **Fig. 2** are presented images with the most complex histological structure, characterizing the tumor state at 7 days after the therapy start. In the histological images, sub-regions corresponding to the most pure areas of the above-mentioned four main morphological constituents of the tumor and the corresponding parts in the OCE-images were selected. In these comparisons, borderlines and transitional tumor zones were excluded. A total of 80 targeted comparisons of tumor zones on the histological images and their corresponding zones on the OCE images were made - these comprise 20 comparisons for each of the four monitored tumor zones (i.e., regions of viable, distrophic, necrotic cells and edema). As a result of the comparisons, a total “stiffness spectrum” of the selected four tumor constituents, for which the averaged stiffness-value histograms (similar to **Fig. 1c5**) were approximated by bell-shape distribution functions that represent their characteristic stiffness values (**Fig. 2f**).

**Fig. 2.**
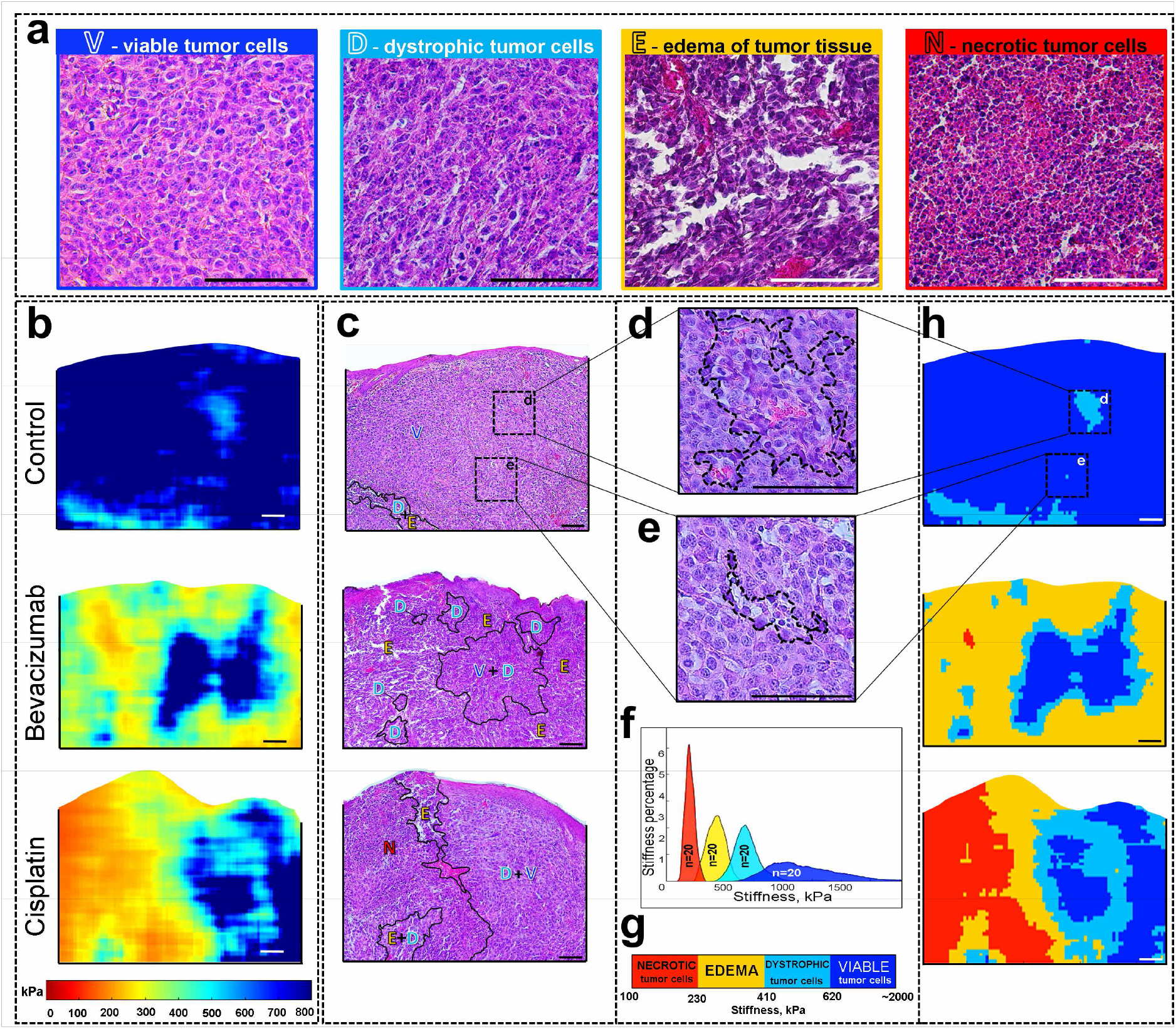
Determination of the specific stiffness ranges for morphological constituents of the tumor. **a**, Zoomed histological images for the four main morphological constituents delineated in the histological sections by an experienced histopathologist. **b-c**, a detailed comparison of the stiffness maps (**b**) derived from OCT scans obtained *in vivo* at day 7 after the initiation of chemotherapy and the corresponding histological images (**c**), in which the black curves show the boundaries of tumor zones identified in the QuPath software (v0.1.2) by a qualified histopathologist without looking at the OCE maps (blind test). **d-e**, Zoomed histology of tumor sites demonstrating high sensitivity of the OCE method in detecting small clusters of cells that are difficult to reveal during the routine histological study. **f**, Total “stiffness spectrum” in which the bell-like functions correspond to average histograms of stiffness values determined for the main morphological constituents. For each of the four morphological constituents, 20 selected tumor zones similar to the representative histology fragments in panel (**a**) were examined. **g**, The graph showing the boundaries of the stiffness ranges and the color palette used for segmentation of the OCE-based stiffness maps shown in (**b**). **h**, Segmented OCE images derived from the stiffness maps (**b**) using the color palette and the characteristic stiffness ranges shown in (**g**). Scale bar for all images = 100 μm.

The so-found four stiffness-distribution functions (with areas normalized to unity) are well separated with small overlaps of no more than 10% of the area between adjacent well-marked peaks (for which the mean stiffness values and standard deviations are presented below). For further segmentation, the stiffness values at the overlap points of the approximating bell-shape functions were selected as the boundaries of the characteristic stiffness ranges (see **Fig. 2g**). As shown in **Fig. 2g**, it was determined that necrotic tumor cells are characterized by the lowest stiffness values from 101 kPa to 230 kPa (average stiffness index of 161 ± 18 kPa); edema has stiffness values from 231 kPa to 410 kPa (average stiffness index of 335 ± 41 kPa); dystrophic of tumor cells has a stiffness range from 411 kPa to 620 kPa (average stiffness of 524 ± 59 kPa). Viable tumor cells has the most rigid structure and is characterized by the highest stiffness values, over 621 kPa, with an average stiffness index of 948 ± 164 kPa (**Fig. 2g**).

The zones corresponding to the so-determined characteristic stiffness ranges for each tumor constituent can be easily automatically indicated in the OCE images, which makes it possible to perform their morphological segmentation automatically. The results of this segmentation are shown in **Fig. 2h**, where the corresponding ranges of stiffness values for each tumor zone are encoded in different colors. Segmented OCE images are much more easily readable due to their higher contrast when compared with the corresponding histological images (**Fig. 2h**).

It is important to note that in **Fig. 2b**, the histological images were marked by the histopathologist in blind recognition, i.e., without looking at the OCE results. Even with a large magnification allowing to see individual cells in the histological section, the histopathologist cannot always find dystrophic cells (denoted as “D” in **Fig. 2**) among viable (denoted as “V”) tumor cells, due to their mosaic distribution. Therefore, a common V + D zone is noted on histological images. However, on the segmented OCE images, the boundaries of these tumor zones are obvious. For example, in the control group, among large fields of viable tumor cells, the histopathologist was not able to isolate small clusters of dystrophic cells. However, in the segmented OCE image, two zones of dystrophic cells are clearly visible. After a more thorough examination of the histological section, these tumor zones were identified as shown in the zoomed fragments of histological section in **Figs. 2d and 2e.** The minimum size of the detected area in Fig. 2e included as few as 10 cells. In this context it can be noted that in comparison with the initial resolution in structural OCT images (~10-15 μm) the distribution of strain in OCE images is spatially smoothed over the window within which the interframe phase gradients (and, therefore, the strains) are estimated. Thus, the expected resolution in OCE images is about 1/2 of the window size corresponding to ~40-50 μm for the OCT scanner used in this study. However, when obtaining the strain/stiffness maps the processing window center is sliding in one-pixel steps (i.e., closer to the resolution scale in the initial structural images). For this reason stiffness inhomogeneities even smaller than the processing-window size can be visualized in segmented OCE images if such a small region exhibits sufficiently high-contrast in stiffness in comparison with the surrounding tissue. Therefore, regions with pronounced contrast in stiffness can be seen in the segmented OCE maps even though the size of these regions (like in **Fig. 2e**) is smaller than the processing-window size.

In **Fig. 2**, it is noteworthy that automatically segmented OCE images and histological images segmented by a histopathologist have a very high degree of similarity, despite the fact that, topographically, the zones of segmented morphological constituents zones may somewhat differ. These differences can be explained by several factors: (1) OCE examination is performed *in vivo* with a slight mechanical compression (and thus deformation) of the tissue, (2) the preparation of histological samples additionally leads to distortion of their shape by osmotic stresses during fixation, dehydration and paraffin embedding, (3) the “same” positions of the OCE scans and histological slices prepared after sacrificing the animal inevitably cannot ideally coincide, whereas neighboring histological sections taken at a distance as small as several tens of micrometers may differ from each other, especially in areas of small-scale structures.

As demonstrates **Fig. 2**, various tumor zones, even very small ones (down to a few-pixels in size), are drawn in detail on the OCE segmented images. These results confirm that the OCE-based examination is exceptionally highly sensitive and even allows for the visualization of such fine variations in the tissue morphology that cannot yet be singled out by a histopathologist in observed sections, because the cells in such places do not yet exhibit clear morphological features allowing their classification as being in one or another morphological state. It is reasonable to conclude that variations in the biomechanical properties of such evolving cells manifest themselves earlier than the visible histological changes. For this reason, OCE can be even more sensitive in revealing early morphological alterations in the tissue than histology.

Thus, it can be reliably said that the OCE method is able very accurately to differentiate between zones of necrotic and viable tumor cells due to the strong separation of their characteristic stiffness ranges that do not have any measurable overlap (see the 1st and 4th peaks in **Fig. 2f**). For adjacent peaks in the stiffness spectrum **Fig. 2f**, the distribution functions of the characteristic stiffness values (**Fig. 2f**) already have some overlap of ~ 10%, which leads to some segmentation uncertainty in regions with stiffness values falling in the overlap zone. Note that this overlap is not exclusively related to the finite accuracy of the measurements, but is mainly caused by the fact that the tumor cells themselves do not instantaneously and simultaneously “switch” from one state to another, so that, for instance, a dystrophic cell may be surrounded by still viable tumor cells and, therefore, the mechanical properties in such zones also do not change abruptly, but gradually evolve within a finite range of stiffness.

The use of higher-resolution OCT systems (for example, similar to**^33–35^** with a cell-level resolution) should make it possible to distinguish the stiffness variations on a smaller scale, down to groups of a few cells or even single histologically changed cells embedded in a fairly homogeneous region of cells of another type; however, typically increase in resolution of OCT systems results in a significantly reduced field of view. In this context it can be emphasized that, for the OCT system used here, with a “typical” resolution of ~10-15 microns in the structural scans, on the one hand, a reasonably wide field of view up to 5-10 μm can be obtained and can readily be extended by stitching together several adjacent OCT scans. On the other hand, such a system already makes it possible to automatically segment OCE scans and visualize different tumor zones comprising fairly small cell agglomerates with sizes of a few tens of microns. For sufficiently high-contrast-in-stiffness regions, this scale can be even less, down to the resolution in the initial structural scans (as in **Fig. 2e** for the small region corresponding to ~10 cells in size, although individual cells are not resolved). It can be pointed out that even if conventional manually performed segmentation of histological sections starts from microscopic consideration of individual cells, the resultant segmentation of these histological sections into macroscopic sub-regions only has an accuracy of tens of microns (and often even lower). Therefore, the result of segmentation based on such conventional histological procedures is comparable with the results of the OCE-based morphological segmentation demonstrated above.

### Correlation between OCE-based segmentation and histological data of tumor response monitoring

Using the above OCE-based technique, we performed monitoring of tumor responses to chemotherapy with parallel verification by histology. As a result of such monitoring, morphologically segmented OCE images were constructed for different time points. For blind verification of the OCE-based segmentation, the corresponding histological images were segmented by an experienced histopathologist without knowing OCE-results. For both types of segmented images (histological and OCE-based), the areas of the revealed tumor zones were calculated and expressed as a percentage of the image area. Morphometric analysis with the calculation of the tumor zones was performed on the corresponding histological images using standard procedures (see Materials and Methods 6-7) and compared with the results of automated calculation of the stiffness-map areas corresponding to each of the determined four characteristic stiffness ranges.

**Fig. 3a** shows a comparison of the areas of segmented tumor zones obtained by the OCE and from the histology. From the bar charts in **Fig. 3a** it is seen that, firstly, there is a high degree of consistency in the percentages of all four main tumor-tissue constituents obtained by the two methods. Secondly, using the OCE, as well as the conventional histological method, one can observe the changes occurring in the tumor in response to chemotherapy. However, using the OCE-segmentation in contrast to standard methods, this can be done non-invasively, *in vivo*, in real time, and on the same animal.

**Fig. 3.**
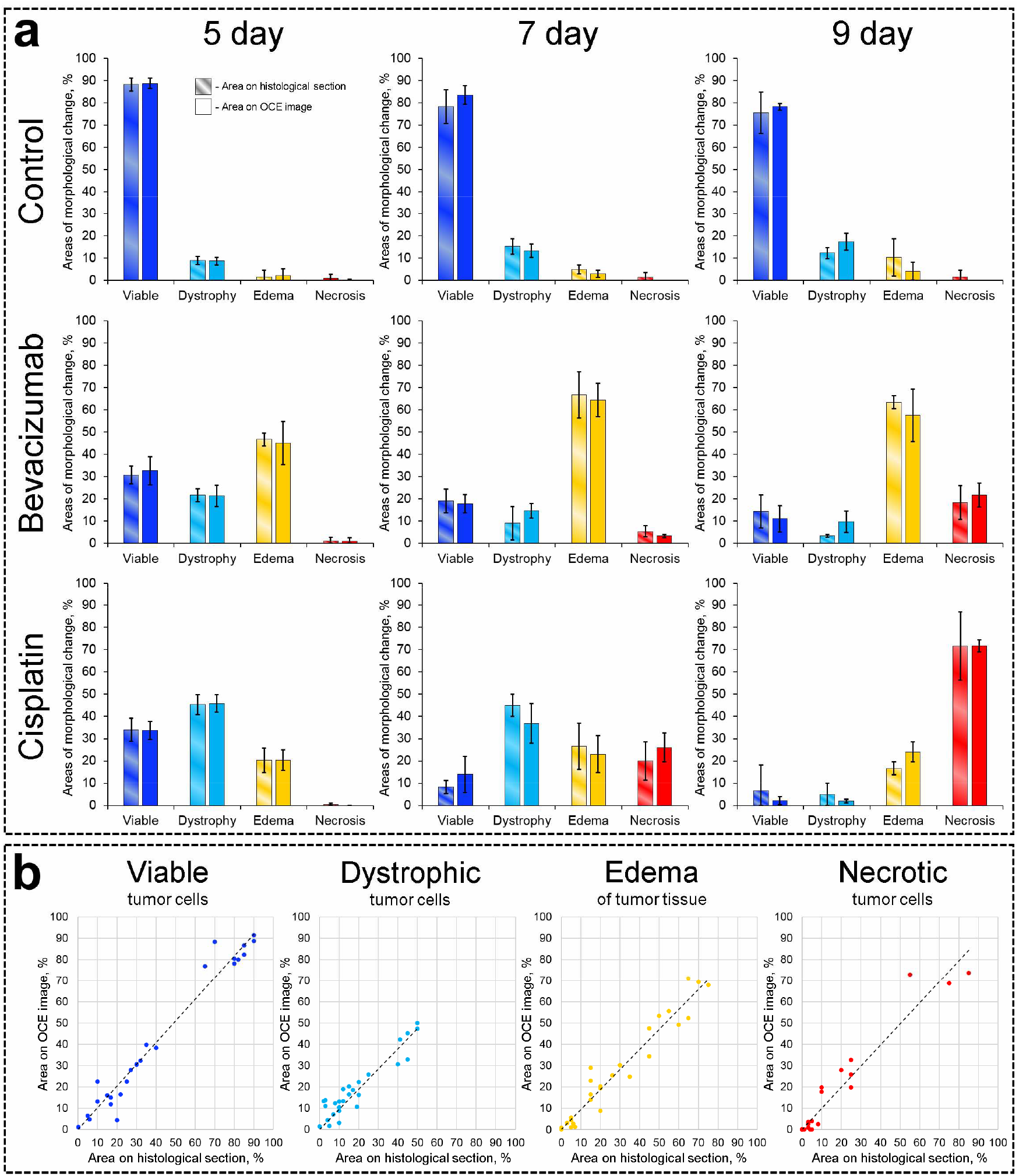
Quantitative comparison of histological and OCE-based segmentation results. **a**, Comparison of the results of histological examination (striped columns) and OCE monitoring (monotone columns). **b**, Relationship between the amount of space occupied by each allocated tumor zone and the percentage of the areas on the OCE images with a stiffness value in the range of this tumor zone. A strong and direct correlation is visible (Pearson correlation coefficient for viable tumor cells r = 0.98, for dystrophic tumor cells r = 0.94, for edema r = 0.97, for necrotic tumor cells r = 0.97).

In addition to the very close results of the two methods of monitoring the development of the tumors (**Fig. 3a**), a high correlation was revealed between the area of each tumor zone highlighted by the histopathologist in the routine way on the histological section and those segmented on the OCE image according to the corresponding elastic modulus range. For all four discussed tumor zones (viable tumor cells, dystrophic tumor cells, edema and necrotic tumor cells), the Pearson correlation coefficient between the areas occupied by them on the histological images and the segmented OCE image is strikingly high, r=0.94-0.98 (see **Fig. 3b**). Thus, the results of morphological segmentation of OCE scans based on monitoring of tumor stiffness are highly consistent with the results of histological examination, which confirms the possibility for using OCE to determine the values of specific stiffness ranges corresponding to various tumor-tissue constituents.

## Discussion

In a broad sense the reported results relate to a vast number of studies aimed at finding methods that accelerate**^36^** / simplify**^37^** / eliminate**^36,37^** any stage of histological examination, or are a convenient alternative to histological study**^38,39^**. Such methods can reduce the cash cost of reagents and save research time. For example, the combination of Raman spectroscopy and autofluorescence in *ex vivo* diagnosis of human skin cancer samples**^38^** has made it possible to differentiate melanoma from basal cell carcinoma with high accuracy, which is a convenient and quick alternative to histological examination. The use of multispectral Muller polarimetry in the *ex vivo* characterization of human colon cancer**^39^** has made it possible to obtain information on the tissue structure without using routine histological examination.

By using combination of third-harmonic generation methods and three-photon fluorescence microscopy, digitally stained multimodal imaging was recently demonstrated. Color-remapped images mimic H&E staining. The method eliminates the stage of special long-term and expensive staining**^37^**. Other studies demonstrate the prospect of using a specially trained neural network to generate a morphological conclusion**^6,36^**, which makes it possible not to have to resort to the help of a qualified histopathologist and to accelerate the pace of the processing of histological material. In addition, our research team is investigating the possibilities of a cross-polarization OCT method in determining the morphology and boundaries of brain tumors**^5^**. These studies illustrate the capabilities of optical methods (including using OCT) for producing high-resolution visualization of tissue structure in places where the use of biopsy studies is completely impossible**^40^**.

Compared with conventional histological examination, the proposed OCE method demonstrates a number of advantages. This method does not require the use of endogenous agents (reagents) and dyes, can be applied *in vivo* or to freshly excised tissue samples**^16,22,41^**. OCE study allows information to be obtained about the tissue under investigation within a few minutes (including recording a signal, processing to obtain a segmented OCE image; it is possible to develop realtime OCE, which will even more significantly reduce the time spent on obtaining the results of the study). By contrast, a routine histological examination takes 3 to 7 days**^42,43^**. It is important to note that there are clinical situations requiring intraoperative, urgent morphological diagnosis. For example, determining the boundaries of tumor resection, establishing the histogenesis of neoplasms, etc. In such cases, an urgent biopsy is performed with the preparation of cryosections. This technique is characterized by fast tissue freezing, bypassing the stage of formalin-fixation and paraffin embedded section prepare procedure. A frozen histological examination takes about 30 minutes from the time the material is taken to the moment the result of the study is obtained, during which time the patient is on the operating table. Moreover, it is known that many studies indicate the significant percentage of false results obtained with such a technique**^44^**. A further development of the OCE method presented here would open up the possibility of real-time implementation of *in vivo* diagnostics of the histological structure of the tissue and its morphological alterations/reactions to therapies.

In this context it can be mentioned that ultrasound shear wave elastography (SWE) was applied in**^45^** to assess the tumor stiffness during thermal therapy to predict the tumor-responders. As the result, SWE mapping and quantification of tumor stiffness was carried out. However, in comparison with OCE ultrasound elastography has significantly lower resolution insufficient for morphological segmentation on a scale comparable with morphometry of histological images.

It can also be noted that alternative histological methods proposed in the literature, using optical methods, are often aimed only at isolating one characteristic component (such as elastin or collagen fibers, or the identification of tumor / non-tumor zones). In this context, OCE opens up the possibility of segmenting several histological components at once. In this paper, four typical tumor zones are considered: necrotic tumor cells, edema, viable tumor cells, and dystrophic tumor cells. Moreover, with a similar approach in the recent study**^16^** on patients with a tumor on the mammary gland made it possible to isolate fibrosis, hyalinosis, inflammatory infiltration and unchanged breast tissue, as well as tumor growth, infiltration of the mammary ducts, and to distinguish between aggregations of tumor cells of different degrees of density. Such results allow us to consider OCE as providing a method for visualizing very fine structures and processes inside highly organized tissues of the human body**^16,41^**.

In addition, the OCE method expands the possibilities of histological examination, enabling the recognition of areas in the tissue with changes that may be missed during routine microscopic examination. Thus, a more complete assessment of the tumor response and thereby the effectiveness of treatment can be achieved.

Knowing both the advantages and the limitations (depth of scanning) of the OCE method, it can be concluded that the method can be valuable in the study of surface structures (skin and mucous, where a biopsy is limited or not advisable) and the effects of various destructive or therapeutic / cosmetic agents on them. OCE can be successfully applied intraoperatively as an express method for determining the morphology and characteristics of tissues, the results of exposure to various therapeutic agents, for determining the boundaries of resection, etc.**^16,20^**. It is very likely that OCE will find uses in oncology as a method for accurately determining the boundaries not only for “healthy tissue/tumor”**^20,46,47^**, but also for “tumor/necrosis” and “healthy tissue/necrosis” boundaries, including the ability to determine other structural features of the tissue.

Besides encouraging results on application of the OCE-based “stiffness-spectrum” approach to assessment/classification of freshly excised breast-cancer samples in**^16^** and the above described *in vivo* results for 4T1 tumor model, this approach was successfully applied in our group to monitor chemotherapy effects on another murine cancer model CT-26, which additionally corroborates possibilities of the proposed OCE-based segmentation to monitor *in vivo* morphological alterations in the same animals in preclinical studies without removing them from the experiment at different time points.

## Conclusion

Possibilities to perform histology-like accurate, nearly real-time morphological segmentation based on optical coherence elastography were demonstrated on a 4T1 breast cancer tumor model treated with two chemotherapeutic drugs with different mechanisms of action. For the first time, by rigorous comparison of OCE images and their corresponding histological images, the stiffness ranges of different tumor zones such as necrotic tumor cells, edema, aggregates of dystrophic tumor cells and viable tumor cells were obtained. The results of the OCE-based morphological segmentation performed *in vivo* were highly correlated with histological images in terms of percentage of the segmented tumor zones. In this regard, the OCE method opens unavailable previously possibilities for assessing the state of tumor either using freshly excised samples or even in vivo to monitor morphological changes during treatment. Numerous applications in other experimental/clinical areas requiring rapid quantitative assessment of tissue morphology can be foreseen.

## Materials and Methods

### 1. Multimodal OCT setup

The elasticity characteristics of tissues were measured using a spectral domain multimodal OCT system (custom-made at the Institute of Applied Physics of the Russian Academy of Sciences) in the compression-elastography mode. The setup has the central wavelength of 1.3 μm and spectral width 90nm, with a power of about 15 mW and can provide 20,000 A-scans per second**^48,49^**. The device had a spatial resolution of 15 μm laterally and about 10 μm (in air) in depth **^50^** in structural images. It is also able to perform real-time angiographic imaging**^7,8,51,52^** and cross-polarization imaging^5,40^.

### 2. OCE imaging

The compression OCE mode is based on estimating the gradients of the interframe variation in the signal phases of adjacent OCT scans during tissue deformation**^9^**. To estimate strains and stiffness, a phase-sensitive technique described in**^11,14,53^** was used together with a robust vector method**^12,13^** for estimating gradients of interframe phase variations that are proportional to interframe strains. Details of obtaining OCE images can be found in**^12,13,15,53^**. For the Young’s-modulus quantification, a calibration (silicone) layer (**Fig. 1b**) with a known stiffness was placed on the tumor surface (**Fig. 1a**). The silicone used here had a Young’s modulus is about 100 kPa as being the most suitable for a study of tumor characterizing changes in stiffness in the range from 50 kPa to 1200 kPa or more. During recording, OCT probe slightly compressed the reference silicone and the underlying tissue.

**Fig. 1c** schematically shows a typical interframe phase difference in the reference silicon and the underlying tissue, for which the interframe strains were found by estimating the axial gradients of the interframe phase variation with averaging over a small rectangular sliding window. The size of this window determined the resolution of the resultant strain/stiffness maps ~40-50 μm.

The silicone layer served as a stress-sensor and thus allowed reconstruction of the stress-strain relationship for the tissue at every point over the OCE-image in a similar manner to**^16^**. Obtaining of stress-strain curves based on finding cumulative strains for several tens of recorded OCT images is described in detail in**^15^**. For real compressed samples, usually strain (and, therefore, stress) inside the homogeneous reference silicone demonstrate pronounced inhomogeneity across the OCE image as shown in **Fig. 4a** for gradually increasing cumulative strains. This inhomogeneous stress in homogeneous silicone is caused by mechanical heterogeneity of the underlying tissue and its non-planar boundary with the silicone and means that pronouncedly different stress is applied to different areas of the tissue within the same OCE frame. This fact should be taken into account, because stiffness of real tissues is pronouncedly stress-dependent as illustrated in **Fig. 4c** by representative stress-strain curves for two different regions within the OCE scan (one for normal tissue and the other for tumor). Both curves are pronouncedly nonlinear with strongly varying local slope that is equal to the local Young modulus (stiffness). Consequently, if the stress applied to the tissue is not specified, interpretation of the observed “stiffness” makes no sense. To exclude this ambiguity in the initial OCE maps with inhomogeneous stress (like in **Fig. 4a**), we construct synthesized (reassembled) maps of cumulative strains corresponding to the same pre-selected stress (and strain) in the reference silicone over the entire synthesized OCE image as shown in Fig. 4b for three pre-selected stress values 3 kPa, 4 kPa and 5 kPa. Such a single reassembled strain map for a pre-selected standardized stress is composed by stitching vertical columns taken from OCE strain-map frames with different numbers in the initially obtained series of OCE maps similar to those in **Fig. 4a**. For such synthesized maps as in **Fig. 4b**, every vertical column corresponds to the same preselected stress (and thus cumulative strain) in the reference silicone, this stress being applied to the underlying tissue. In such a way we ensure that, for every position on the OCE map, we obtain the stress and strain values corresponding to stresses 4±1 kPa on the nonlinear “stress-strain” curves like those in Fig. 4c. Then, for the central stress value 4 kPa, the local slope of the “stress-strain” curve (i.e., the Young modulus for 4 kPa) is approximated by the chord connecting the pre-selected stress points (say, 4-1=3 kPa and 4+1=5 kPa as in Fig. 4c). The chosen central stress (here, 4 kPa was empirically chosen) can be adjusted for a particular heterogeneous tissue to obtain strains in the examined region from fractions of one percent to several percent (that do not yet damage the tissue and are fairly easily measured). For such a method of stiffness mapping, the ambiguity due to the tissue nonlinearity is excluded and the result is much more robust in comparison with straightforward estimates based on interframe stress and strain increments. The so-constructed map of the Young modulus standardized for 4 kPa stress over the entire OCE scan is shown in **Fig. 4e**.

**Fig. 4.**
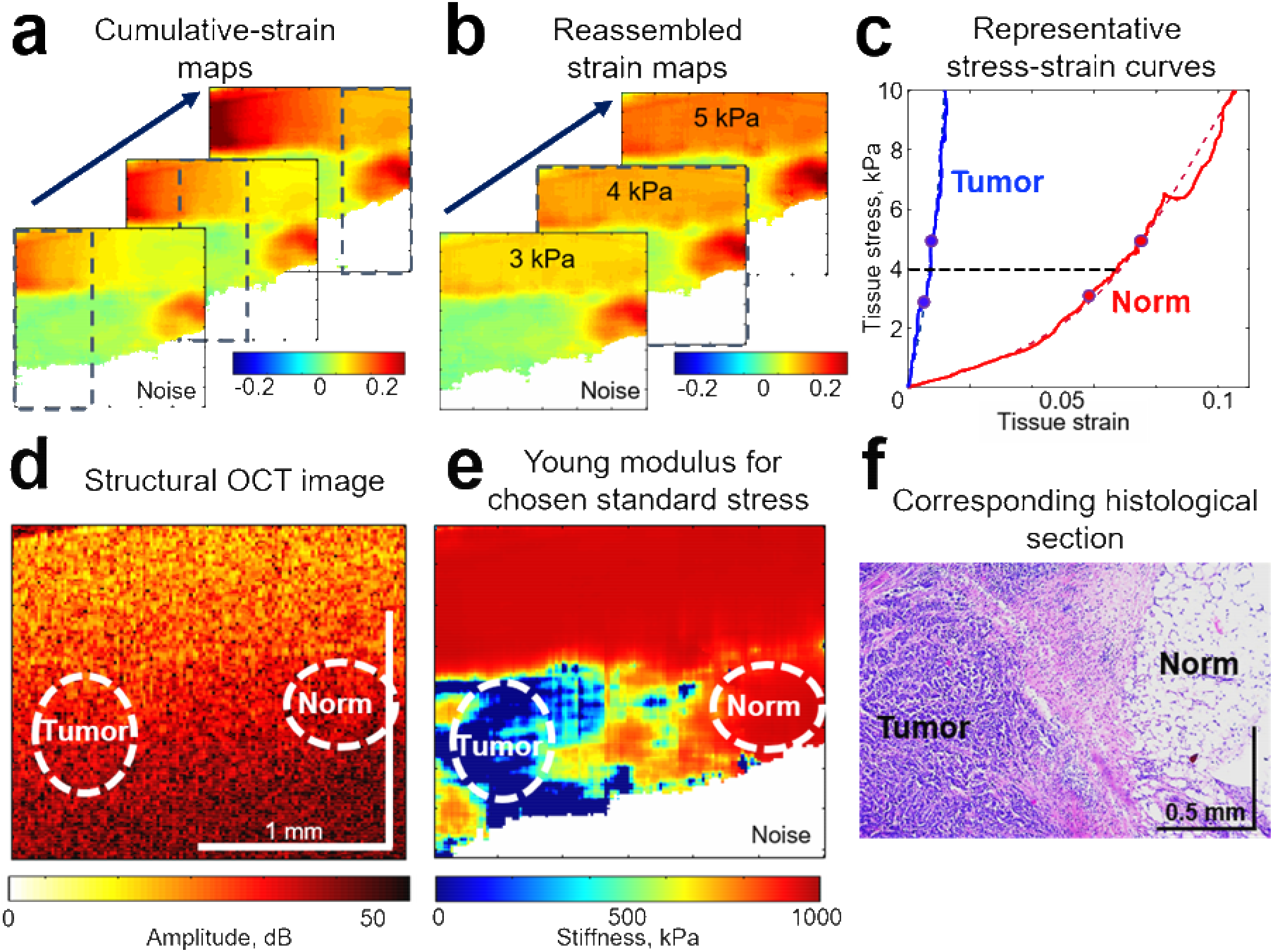
Schematically shown experimental OCE procedures for obtaining synthesized stiffness maps ensuring standardized pressure exerted on structurally/geometrically inhomogeneous tissue. **a**, A series of initially calculated maps of cumulative strains that are essentially inhomogeneous within the reference silicone layer. **b**, Reassembled (synthesized) maps of cumulative strains corresponding to the three pre-selected levels of pressure (3, 4 and 5 kPa) in the reference silicone layer. **c**, Representative nonlinear stress-strain curves obtained for two regions essentially differering in stiffness (normal and tumorous region). The curves demonstrate that the local slope of the stress-strain curves (i.e. the Young’s modulus) may change several times even for apparently moderate strain ~several per cent. **d**, Typical initial structural OCT scan. **e**, Synthesized Young’s-modulus map corresponding to the standardized stress of 4 kPa in the reference silicone across the entire scan. **f**, histological section corresponding to the same position.

The derived stiffness maps (like in **Fig. 4e**) could be superimposed with conventional structural OCT images (**Fig. 4d**) and compared with the corresponding histological slices to find characteristic stiffness ranges (“stiffness spectra”) for various tumor zones as described in detail in the discussion of **Fig 2**. Then the stiffness maps (like those in **Fig. 2c**) can be automatically segmented into regions corresponding to the preliminary found stiffness ranges (see **Fig. 2h**) and in any chosen regions of interest percentages of pixels (percentages of the analyzed area) corresponding to various ranges of stiffness could be automatically calculated and used to monitor variations in the tissue morphology (as shown in **Fig. 3a**). Also correlation between the results of OCE-based segmentation and morphological segmentation of conventional histological images could be found for the four tumor zones of primary interest: regions of viable tumor cells, dystrophic tumor cells, tumor edema, and necrotic tumor cells (see **Fig. 3b**).

### 3. Experimental animals and cell lines

Balb/c female mice (6–8 week old, n=36 divided in 3 equal groups) were used in the experiments performed in accordance with the European Convention for the Protection of Vertebrate Animals used for Experimental and Other Scientific Purposes (ETS 123), the experimental protocol being approved by the Research Ethics Board of the Privolzhsky Research Medical University (REB#2, granted January 29, 2018). A mouse mammary carcinoma 4T1 cell line was cultured in RPMI-1640 with 10% FCS, 1% glutamine, 10 units/mL penicillin and 10 μg/mL streptomycin and maintained in an incubator at 37 °C in an atmosphere of 5% CO2. The cells were subcultivated 3 times at 80% of confluence before injection. To obtain the animal tumor model, the cells were harvested using trypsin-EDTA solution, counted and resuspended in phosphate buffered saline (PBS). The cells were inoculated subcutaneously in auricle tissue of the Balb/c mice at a concentration 2×10^5^ cells per 20 μl PBS**^54^**. The relatively small tumor sizes enable accurate 3D size determination; this is very important to calculate tumor volume accurately. The histological structure and other details of this tumor model are described in**^55^**. As shown earlier**^56,57^**, the 4T1 model is a triple negative tumor, morphologically similar to duct breast cancer and characterized by a high degree of malignancy and high metastatic activity. It is also important that the surface location of the tumor allows for visual assessment of its growth and monitoring of the changes in the tissue elasticity using OCE**^22^**.

### 4. Chemotherapy drugs

A comparative study of the action of two types of chemotherapeutic drugs was carried out. Bevacizumab belongs to the angiogenesis inhibitor family: it is a monoclonal antibody that targets vascular endothelial growth factor (VEGF-A). VEGF-A is a growth factor protein that stimulates angiogenesis in a variety of diseases, especially in cancer**^58^**. Inhibition of VEGF by Bevacizumab affects tumor growth by several mechanisms, including the inhibition of new vessel growth and induction of endothelial cell apoptosis. Moreover, it affects vessel function by vasoconstriction and vessel normalization**^59^**.

Cisplatin belongs to the platinum-based antineoplastic family of drugs used for cytostatic antitumor chemotherapy. It works, in part, by binding to DNA and inhibiting its replication**^30^**. By interfering with DNA replication Cisplatin kills the fastest proliferating cells, which, in theory, are the carcinogenic ones**^60,61^**.

Treatments started 2 days after tumor cell inoculation. Bevacizumab and Cisplatin were administrated in an intraperitoneal manner, in doses of 15 and 6 mg/kg respectively, according to a previously proposed treatment regimen**^62,63^**. Five cycles were administered over 2 weeks, three times per week (at days 0, 2, 5, 7 and 9).

### 5. Macroscopic evaluation of tumors

At each time point (days 0, 2, 5, 7, 9, 12), photographs of the tumors were taken on a stereomicroscope (Axio Zoom v16 ZEISS). During the first 12 days, the therapeutic effect was assessed by following the tumor growth kinetics in the control and therapeutic groups. Tumor growth and progression were monitored by measurement of the tumor size with a caliper device three times per week. Tumor volume (V, mm^3^) was determined by the equation**^64^**:

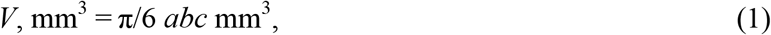

where *a* is the length of the tumor, *b* is its width and *c* is the height. Data was plotted as the average tumor size for the 10 mice in each treatment group. Since at the baseline (day 0), the tumor volume varied greatly from mouse to mouse, the values of tumor volume on days 1–12 were normalized per the baseline value to obtain the ratio *V/V*0 for the very same mouse. The tumor growth inhibition coefficient (TGI) was calculated on day 12 from the beginning of treatment using the formula**^65^**:

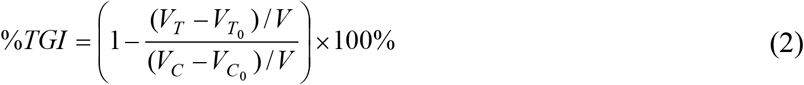

where *V_T_* is the mean tumor volume on day 12 after the start of chemotherapy, *V*_*T*0_ is the mean tumor volume on the day of the start of chemotherapy; *V_C_* and *V*_*C*0_ are the corresponding means for the control group. This derived metric quantifies the efficacy of the treatment assessed volumetrically while accounting for volume changes due to natural disease progression over the monitoring time interval. TGI > 100% means that the tumor volume decreased after chemotherapy (high efficacy of treatment); 100% > TGI > 50% means that the tumor volume moderately increased (medium efficacy of treatment), TGI < 50% tumor volume greatly increased (ineffective treatment)**^66^**.

### 6. Histological examination

Animals were sacrificed by dislocating the cervical vertebrae under anesthesia. The excised tumors were embedded in paraffin. Several cross sections were made from the center of the tumor. Histological sections were stained with hematoxylin and eosin (H&E). The histopathological study included the identification of viable tumor cells, necrotic tumor cells, edema and dystrophic tumor cells (atypical / pathological mitoses, cells with karyopicnosis of the nucleus, karyorexis of the nucleus, karyolysis of the nucleus, nucleus vacuoles and cytoplasmic vacuoles).

Quantification of these tumor zones was carried out in QuPath image analysis software (v0.1.2). Using this software, a histopathologist identified the above mentioned tumor zones and their boundaries. The percentage of the area occupied by each tumor zones was equal to the ratio of the sum of all the areas of the given structure on the histological image to the total image area (the area corresponding to the size of the OCE images taken under the histological mark). For accurate comparison of the histological images with the OCE images, histological paint had been placed at the center of the tumor. For analysis, one section was taken from each tumor.

### 7. Statistical analysis

Statistical analysis was performed in Statistica 10.0 software. Pearson’s correlation coefficient was used for comparison between areas of various tumor zones determined by histology and from the automatic segmentation of OCE images. Statistical analysis of tumor growth was performed using the one-tailed Student’s t-test. P values less than 0.05 were considered statistically significant.

## Data availability

The data that support the findings of this study are available from the corresponding author upon request.

## Acknowledgments

Development of an approach to quantitative assessment of OCE stiffness maps and determining the characteristic tissue stiffness ranges was supported by the RSF grant 18-75-10068. The method of obtaining synthesized stiffness maps for a standardized pressure was developed under the support of RFBR grant 18-32-20056. Development of approach to assessment of tumor response to chemotherapy was funded by RFBR, project number 19-315-90087. The authors are grateful to Prof. Alex Vitkin (University of Toronto) for valuable discussions.

## Author contributions

N.D.G., V.Y.Z. and M.A.S. designed the study. A.A.P., A.A.S. and E.V.G. collected the experimental data. OCE imaging method was developed by V.Y.Z., A.A.S., A.L.M. and L.A.M. The results were interpreted by M.A.S., A.L.M., A.A.S., A.A.P., V.Y.Z. and S.S.K. A.A.P., M.A.S. and V.Y.Z. wrote the manuscript. N.D.G., E.V.G. and E.V.Z. edited the manuscript. A.A.P., S.S.K., M.A.S. and V.Y.Z. prepared the figures. All authors reviewed the manuscript.

